# A comparative analysis of toxin gene families across diverse sea anemone species

**DOI:** 10.1101/2024.12.13.628455

**Authors:** Hayden L. Smith, Daniel A. Broszczak, Chloe A. van der Burg, Joachim M. Surm, Libby Liggins, Raymond S. Norton, Peter J. Prentis

## Abstract

All species from order Actiniaria (sea anemones) are venomous, even though most are of no threat to humans. Currently, we know very little about the toxin gene complement of highly venomous members of this order. To address this gap in knowledge, we sequenced the transcriptome of the highly venomous and medically significant Hell’s Fire sea anemone, *Actinodendron plumosum*, as well as five distantly related species, *Cryptodendrum adhaesivum*, *Epiactis australiensis*, *Heteractis aurora*, *Isactinia olivacea* and *Stichodactyla mertensii*. We used bioinformatic approaches to identify their toxin gene complements and performed a comparative evolutionary analysis of seven understudied toxin families. Of the 16 toxin families identified, 12-40 candidate toxins were found in the six new sea anemone transcriptomes, with only 12 candidates in eight toxin families identified in *A. plumosum*. Across 26 sea anemone species, six neurotoxin families showed evidence of taxonomic restriction, whereas the phospholipase A2 toxin family was ubiquitously distributed. Additionally, we identified two alternative forms for the phospholipase A2 toxin family, a 10- and 14-cysteine framework, which warrant further structural and functional characterisation. Overall, we have identified a comprehensive list of toxins from a wide diversity of sea anemone species that provides the basis for future research to structurally and functionally characterise novel candidates for use as therapeutics or for agricultural applications.

## 1. Introduction

Venoms and toxins from reptiles, cone snails and arachnids have been well characterised, because of their importance to the pharmaceutical industry and ease of collection (Israel et al., 2017; King & Hardy, 2013; Lewis et al., 2012; Pennington et al., 2018). This research has led to breakthrough discoveries in the application of these proteins and peptides in a range of fields, such as pharmacology (Israel et al., 2017; King & Hardy, 2013; Lewis et al., 2012) and drug discovery (de Oliveira Amaral et al., 2019; Harvey, 2014; Muttenthaler et al., 2021; Pennington et al., 2018; Saez & Herzig, 2019). In contrast, venoms from phylum Cnidaria have been largely understudied, with relatively few species examined for the applications of their peptide toxins in the biomedical and agricultural industries (Bosmans & Tytgat, 2007; King, 2011; Mariottini, 2016; Yan et al., 2014). This is quite surprising given the diverse applications of the few candidates that have been examined, such as ShK, APETx4, ATX-I, ATX-II and ShI (Beeton et al., 2011; Chi et al., 2012; Honma & Shiomi, 2006; Kem et al., 1989; Moreels et al., 2017; Norton, 2009; Pennington et al., 2009; Prentis et al., 2018; Schweitz et al., 1981; Sintsova et al., 2023; Tarcha et al. 2017; Upadhyay et al., 2013). The importance of these medically significant toxins highlights the need for more research to examine toxin repertoires in other cnidarian species.

Within phylum Cnidaria, sea anemones (order Actiniaria) represent a particularly rich and structurally diverse source of peptide toxins (Elnahriry et al., 2019; Madio et al., 2019), which they use for a wide range of ecological purposes, including intra-specific combat, prey capture, defence against predators and digestion (Ashwood et al., 2020; Schendel et al., 2019; Surm et al., 2024a). Many of these toxins are encoded by genes from large gene families, such as the potassium channel toxin (Castañeda & Harvey, 2009; Jouiaei et al., 2015), sodium channel toxin (Jouiaei et al., 2015; Moran et al., 2009), and actinoporin (Macrander & Daly, 2016; Valle et al., 2015) superfamilies. Comparative studies for toxin gene families from species of medical significance have not been conducted in detail; instead there has been a focus on specific species and genera, such as *Nematostella vectensis* (Columbus-Shenkar et al., 2018; Moran et al., 2008; Moran et al., 2013; Sachkova et al., 2019), *Stichodactyla* spp. (Madio et al., 2017, Rivera-de-Torre et al., 2018) and *Actinia* spp. (Moran et al., 2008; Surm et al., 2019a; Surm et al., 2019b). One medically significant species, *Actinodendron plumosum*, commonly known as the Hell’s Fire sea anemone, has a severe and painful sting with some symptoms including burning, necrosis, ulceration, and fever (Haddad et al., 2009; Hansen & Halstead, 1971). With the exception of a few species (Ashwood et al., 2021; Macrander et al., 2016; Uechi et al., 2010), the gene family complements of most highly venomous and painful sea anemone lineages have not been studied, despite their potential importance in pharmacology and drug discovery (Pennington et al., 2018; Prentis et al., 2018).

In sea anemones, genes encoding peptide and protein toxins from various neurotoxin families and the phospholipase A2 family have received only scant attention, with no detailed analysis of gene family evolution across taxonomic families. Consequently, new gene variants of known sea anemone toxin families may be found and prove to be a valuable resource for future biodiscovery efforts. We have therefore undertaken deep transcriptome sequencing to determine the diversity of toxin genes in six sea anemone species from five different taxonomic families (Actinodendridae, Thalassianthidae, Actiniidae, Heteractidae, Stichodactylidae) and compared these to toxin genes identified in transcriptomes from closely and distantly related sea anemone species. Using this sequence information, we then examined the diversity and evolution of seven toxin families that have not been previously investigated in detail; acrorhagin type I (AcrI), CjTL, inhibitor cystine knot (ICK), potassium channel toxin (KTx) type V, phospholipase A2 (PLA2), small cysteine-rich protein (SCRiP) and sodium channel toxin (NaTx) type III.

## 2. Materials and Methods

### 2.1 Sampling, RNA extraction and library preparation

*Actinodendron plumosum*, *Cryptodendrum adhaesivum*, *Heteractis aurora* and *Stichodactyla mertensii* were obtained from Cairns Marine (Queensland, Australia). *Epiactis australiensis* was collected from tidal rock pools at Point Arkwright (Queensland, Australia). *Isactinia olivacea* was collected from Hatfields Beach (Auckland, New Zealand) and stored in RNAlater. Species were identified based on morphology and alignment of transcripts to ribosomal genes from the NCBI database. All sea anemones, except *A. plumosum* and *I. olivacea*, were acclimated to a recirculating tank system and fed prawn every three days for one week at QUT (Queensland, Australia) prior to RNA extraction. Due to its high toxicity, *A. plumosum* was flash frozen after transport from Cairns Marine and stored in -80°C freezer until RNA extraction. Total RNA was extracted from whole organism using a previously optimised extraction protocol for marine invertebrates (Prentis & Pavasovic, 2014). RNA quality and quantity were determined using an RNA chip on the Bioanalyzer 2100 (Agilent Technologies). High-quality RNA samples were used for TruSeq Stranded RNA Library Preparation and sequenced using 150bp paired-end chemistry on a HiSeq 2500 at Novogene (China).

### 2.2 Transcriptome assembly and contaminant removal

Raw sequence data were quality checked and assembled using the Trinity *de novo* assembler software (v2.13; Haas et al., 2013). Only high quality reads (> Q30, < 1% ambiguities) were used for assembly using default settings, with the trimmomatic flag used to remove any low quality reads and adaptors as well as the stranded option (Haas et al., 2013). All raw reads were submitted to the NCBI Sequence Read Archive (SRA) under BioProject PRJNA507679. Potential symbiont contamination was removed from transcriptomes using the Psytrans Python script (https://github.com/sylvainforet/psytrans). Symbiont reference sequences were combined from the genome protein models of *Symbiodinium microadriaticum*, *S. minutum* and *S. kawagutii* (Aranda et al., 2016; Lin et al., 2015; Shoguchi et al., 2013). Default script settings were used to analyse the sea anemone transcriptomes, except for the following script modifications; maxBestEvalue = 1e-30, numberOfSeq = minimum blast hit count, Training Proportion = 1.0, Line 566: if nSeqs < options.args.numberOfSeq:, Line 572: for i in xrange(trainingMaxIdx, options.args.numberOfSeq).

TrinityStats.pl script from Trinity *de novo* assembler software (v2.13; Haas et al., 2013) was used to determine quality statistics of transcriptome assemblies. BUSCO software (v3.0.1) was used to determine the relative completeness of the transcriptome assemblies using the presence of the 978 metazoan single-copy orthologs (Simão et al., 2015) with a score of > 80% considered high quality.

An additional 20 RNA-seq datasets obtained from NCBI that were previously assembled (Ashwood et al., 2021; Cuttitta et al., 2017; Mitchell et al., 2020; van der Burg et al., 2016; van der Burg et al., 2020; Xu et al., 2022) were used for subsequent toxin candidate identification, and phylogenetic and molecular analyses (Supplementary Materials 1).

### 2.3 Toxin transcript identification

Toxin and toxin-like transcripts were identified using methods published previously (Smith et al., 2023; Smith et al. 2024). Briefly, full-length open reading frames with a minimum of 30 amino acids were predicted using TransDecoder (v3.0.1; https://github.com/TransDecoder/TransDecoder/releases). SignalP (v5.0) was used to identify the presence of a signal peptide, and toxin candidates were identified using BLAST searches on UniProt (https://www.uniprot.org/) against the Swiss-Prot database for cnidarian toxins (Taxonomy: Cnidaria (6073) and Keyword: Toxin (KW-0800) qualifiers). Candidates were manually curated to ensure BLAST hits had similarity to known toxin peptides and proteins with high confidence (P-value ≤ 1.0e-6), specifically removing candidates with incomplete PFAM domain structures based on SMART searches (http://smart.embl.heidelberg.de/) or incorrect cysteine-frameworks based on characterised toxins in the UniProt and hmmscan (https://www.ebi.ac.uk/Tools/hmmer/search/hmmscan) databases (Supplementary Material 1).

### 2.4 Phylogenetic analysis of toxin gene families

Phylogenetic analyses were performed using previously published methods (Smith et al., 2023; Smith et al. 2024; Surm et al., 2019b). Briefly, trees were constructed for seven toxins of interest (acrorhagin type I, CjTL, inhibitor cystine knot, potassium channel toxin type V, phospholipase A2, small cysteine-rich protein and sodium channel toxin type III) due to a lack of comparative evolutionary analyses prior to this study. MUSCLE (v5.1; Edgar, 2004) was used to determine an alignment for each toxin family and a phylogenetic tree was constructed using IQ-TREE (v1.6.12; http://iqtree.cibiv.univie.ac.at/; Trifinopoulos et al., 2016) with 1,000 bootstrap alignments. Okabe-Ito colour-blind barrier free palette (jfly.uni-koeln.de/color) was used to visualise bootstrap support (poor: ● 0-49, low: ● 50-69, moderate: ● 70-79, strong: ● 80-99, very strong: ● 100) in addition to bootstrap values.

## 3. Results

### 3.1 Transcriptome assembly and annotation

Transcriptome assemblies had high quality statistics (N50 > 1,000 and BUSCO Score > 90%) with the exception of *I. olivacea* (Table 1). The *I. olivacea* transcriptome had a relatively low gene count, < 50,000, and the lowest N50 value of 1,380, as well as a lower quality BUSCO score at 77.8%. Statistical metrics were rerun for the raw data (prior to psytrans filtering) of *I. olivacea* which resulted in 86,332 genes, a N50 of 1,139, a BUSCO score of 86.8%, resulting in a 49.5% reduction in genes due to Psytrans filtering. These metrics suggest that the raw data are incomplete for this species, possibly due to storage and transport in RNAlater.

**Table 1:** Summary statistics for transcriptome assembly and completeness (BUSCO) after contamination removal with Psytrans.

| Species | Trinity Stats |  |  |  |  | BUSCO Score % |
| --- | --- | --- | --- | --- | --- | --- |
|  | Genes | Transcripts | GC % | N50 | Median contig |  |
| *Actinodendron plumosum* | 123,166 | 175,633 | 38.22 | 1,563 | 408 | 92.1 |
| *Cryptodendrum adhaesivum* | 77,515 | 91,687 | 39.8 | 1,417 | 425 | 93.7 |
| *Epiactis australiensis* | 62,029 | 92,073 | 39.24 | 1,945 | 593 | 94.2 |
| *Heteractis aurora* | 90,718 | 137,762 | 40.27 | 1,667 | 458 | 92.9 |
| *Isactinia olivacea* | 43,535 | 48,832 | 39.56 | 1,380 | 509 | 77.8 |
| *Stichodactyla mertensii* | 83,154 | 121,429 | 38.9 | 1,714 | 474 | 92.4 |

### 3.2 Toxin transcript identification

Between 12-40 transcripts that encode toxins across 16 toxin protein families known to occur in sea anemones were found in the six new transcriptomes (Table 2). While *A. plumosum* had the most genes, it had the least number of toxin candidates (12) and toxin families (8) compared with the other five species. Conversely, *S. mertensii* had ∼40,000 fewer genes than *A. plumosum* but the most toxin candidates and a greater number of toxin families, 40 and 11, respectively. Identification of known toxins for *A. plumosum*, as well as *E. quadricolor*, revealed a relatively large number of duplicate (identical full ORF) genes and isoforms (Supplementary Material 1). Toxin identification of previously published data for superfamilies Edwardsioidea and Metridioidea showed the lowest counts for the 16 toxin families of interest, specifically between 10-21 and 8-27 candidate toxins, respectively (Supplementary Material 1).

**Table 2:**
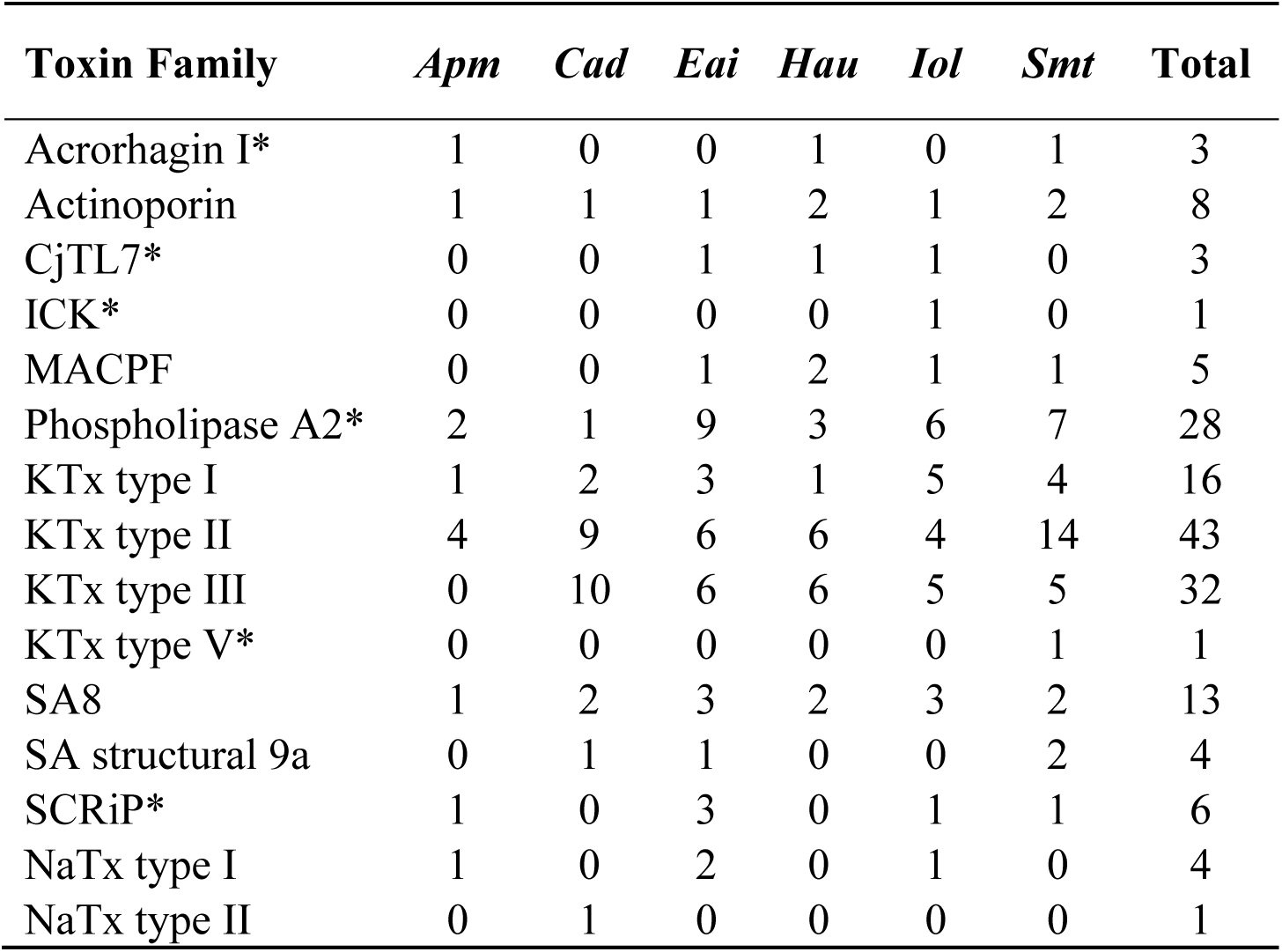
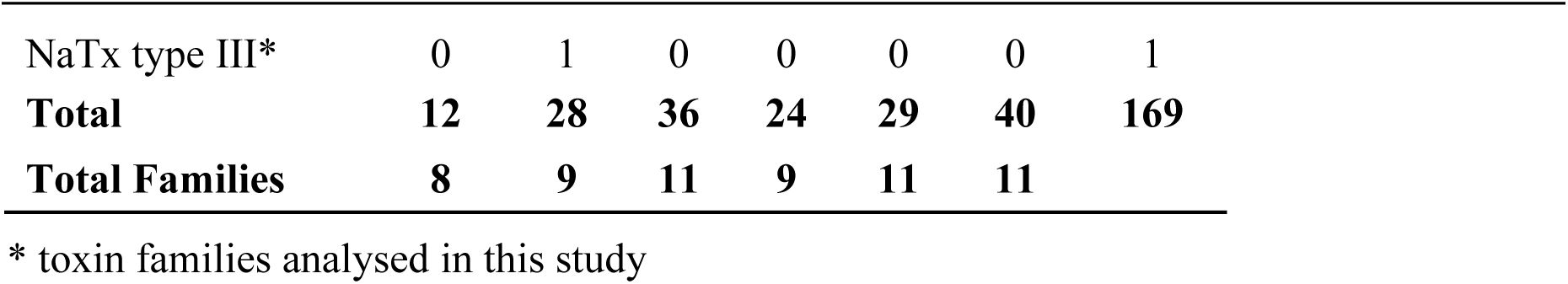
Number of putative toxin-encoding transcripts from the 16 toxin families identified in the six new transcriptomes. Species Name Abbreviations: *A. plumosum* (Apm), *C. adhaesivum* (Cad), *E. australiensis* (Eai), *H. aurora* (Hau), *I. olivacea* (Iol), *S. mertensii* (Smt). Toxin Family Abbreviations: potassium channel toxin (KTx), sea anemone (SA), small cysteine-rich protein (SCRiP), sodium channel toxin

| Toxin Family | *Apm* | *Cad* | *Eai* | *Hau* | *Iol* | *Smt* | Total |
| --- | --- | --- | --- | --- | --- | --- | --- |
| Acrorhagin I* | 1 | 0 | 0 | 1 | 0 | 1 | 3 |
| Actinoporin | 1 | 1 | 1 | 2 | 1 | 2 | 8 |
| CjTL7* | 0 | 0 | 1 | 1 | 1 | 0 | 3 |
| ICK* | 0 | 0 | 0 | 0 | 1 | 0 | 1 |
| MACPF | 0 | 0 | 1 | 2 | 1 | 1 | 5 |
| Phospholipase A2* | 2 | 1 | 9 | 3 | 6 | 7 | 28 |
| KTx type I | 1 | 2 | 3 | 1 | 5 | 4 | 16 |
| KTx type II | 4 | 9 | 6 | 6 | 4 | 14 | 43 |
| KTx type III | 0 | 10 | 6 | 6 | 5 | 5 | 32 |
| KTx type V* | 0 | 0 | 0 | 0 | 0 | 1 | 1 |
| SA8 | 1 | 2 | 3 | 2 | 3 | 2 | 13 |
| SA structural 9a | 0 | 1 | 1 | 0 | 0 | 2 | 4 |
| SCRiP* | 1 | 0 | 3 | 0 | 1 | 1 | 6 |
| NaTx type I | 1 | 0 | 2 | 0 | 1 | 0 | 4 |
| NaTx type II | 0 | 1 | 0 | 0 | 0 | 0 | 1 |
| NaTx type III* | 0 | 1 | 0 | 0 | 0 | 0 | 1 |
| **Total** | **12** | **28** | **36** | **24** | **29** | **40** | **169** |
| **Total Families** | **8** | **9** | **11** | **9** | **11** | **11** |  |

\* toxin families analysed in this study

### 3.3 Toxin complement comparisons to other sea anemone species

Overall, 951 toxin and toxin-like genes were identified across the 26 transcriptomes analysed, including 422 enzymes, 481 neurotoxins and 48 with limited characterisation (Supplementary Material 1). On average, superfamily Actinioidea had a greater number of toxin and toxin-like genes than superfamilies Edwardsioidea and Metridioidea, with approximately 30, 15 and 18, respectively, although, a larger sample size for edwardsioidean and metridioidean species would be needed to further support this result. Five toxin families were present in all six of the new transcriptomes, actinoporin, KTx type I, KTx type II, PLA2 and sea anemone 8. Of these, PLA2 was present across all 26 transcriptomes analysed, whereas, actinoporin, KTx type I, KTx type II and sea anemone 8 were present in 17, 18, 25 and 24 of the 26 transcriptomes analysed, respectively (Supplementary Material 1: “Compare1” tab).

#### 3.3.1 Acrorhagin I

Acrorhagin type I (Honma et al., 2005; Krishnarjuna et al., 2021) toxins were found exclusively in superfamily Actinioidea with a single copy found in 10 species and two copies found in *A. tenebrosa* (Figure 1a). *Actinia tenebrosa* was the only species with the canonical short C-terminal (*A. equina*, Aeq_Q3C257 and Aeq_Q4C258), whereas all other sequences had a very strong conservation of 20-21 amino acids after the canonical stop site. It is possible that post-translational cleavage occurs at the KR (Lys-Arg) site for these longer sequences, resulting in toxins with similar structures and functions to the characterised toxins, Aeq_Q3C257 and Aeq_Q3C258 (Figure 1a). Phylogenetic analysis confirmed the clustering of these two toxins and Ate 1 separate from all other sequences (Figure 1b). Species groupings were congruent with well-known phylogenetic relationships, and strong support was observed for three groups, Apm_1 and Mgr_1, Cad_1 and Hhy_1, and Cgg_1 and Hau_1 (Figure 1b).

**Figure 1:**
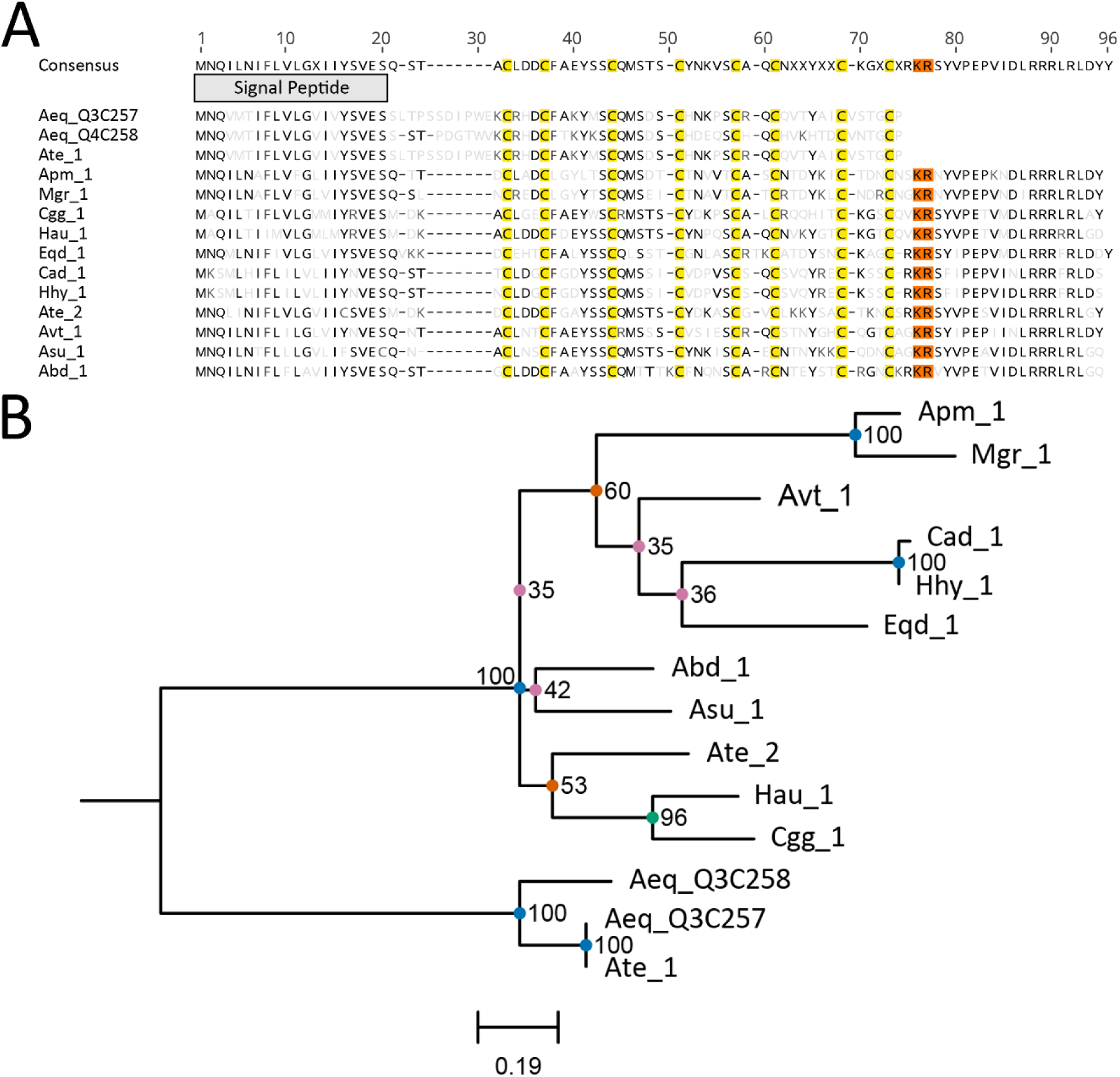
Amino acid sequence alignment and phylogenetic tree for the acrorhagin type I toxin family sequences. (A) Alignment of candidate sequences with highlighted sections for predicted signal peptide, cysteine residues and potential cleavage site motifs. Light grey letters represent non-consensus amino acids. (B) Maximum-likelihood phylogenetic tree of candidate toxins. Support values shown as Maximum Likelihood bootstrap (0-100). Gene transcripts were identified based on their species name followed by count number. Sequences denoted with a six-digit alphanumeric code are from the Swiss-Prot/UniProt databases.

#### 3.3.2 Toxin CjTL

CjTL (Babenko et al., 2017) toxins were found exclusively in superfamily Actinioidea with a single copy found in nine species and two copies found in *A. tenebrosa* (Figure 2a). However, only the Uniprot sequence P0DPE6 and Cgg_1 have a KR cleavage site, which highlights the need for further structural and functional analysis of these candidates to verify toxin activity. Importantly, there is a very strong conservation of amino acids throughout the sequence alignment of all candidate toxins (Figure 2a). Phylogenetic analysis for the CjTL sequences had moderate support with no clear clustering outside of Avt_1 and Iol_1, and Cgg_1, Hau_1 and Hdo_1, which have very strong support (Figure 2b).

**Figure 2:**
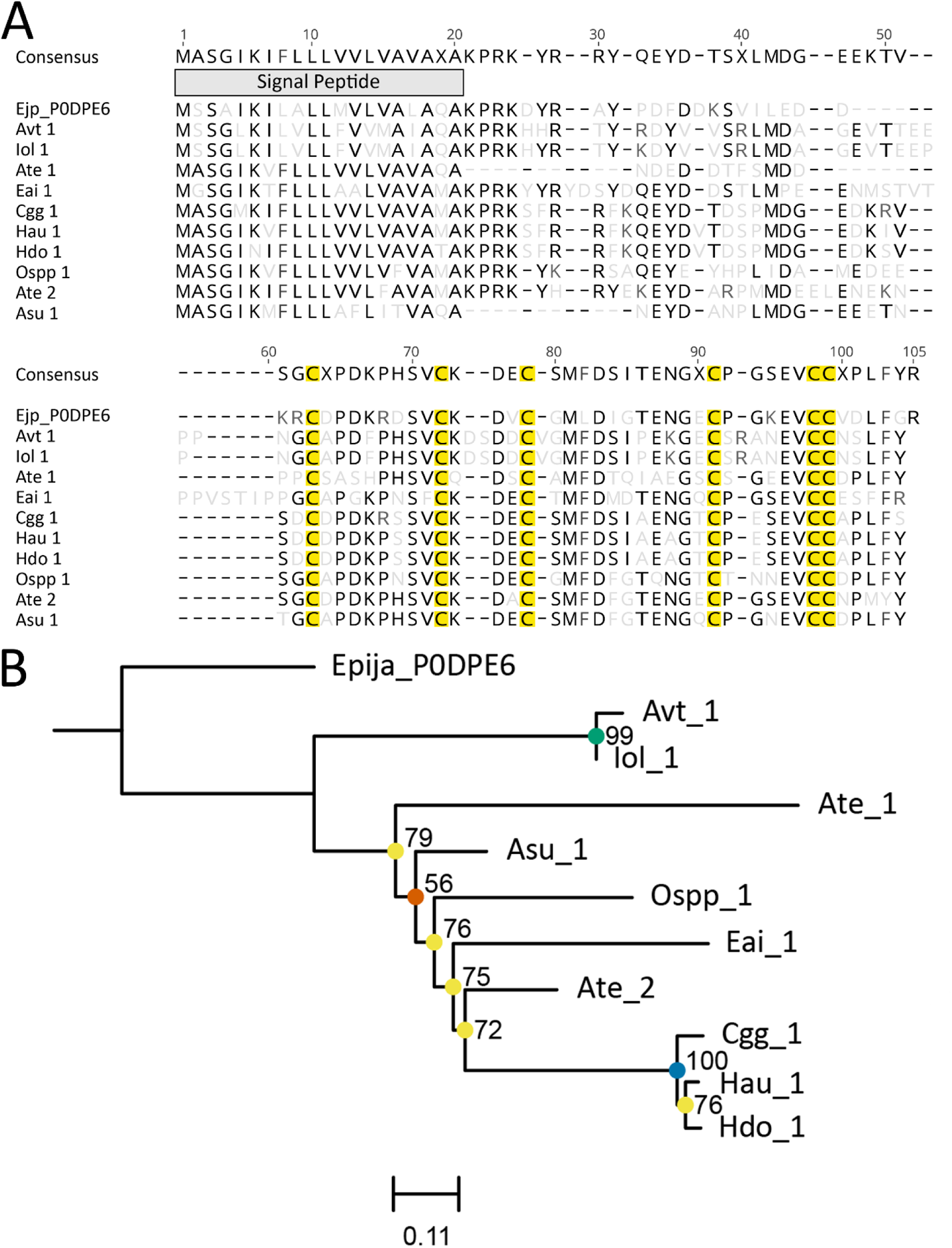
Amino acid sequence alignment and phylogenetic tree for the CjTL toxin family sequences. (A) Alignment of candidate sequences with highlighted sections for predicted signal peptide and cysteine residues. Light grey letters represent non-consensus amino acids. (B) Maximum-likelihood phylogenetic tree of candidate toxins. Support values shown as Maximum Likelihood bootstrap (0-100). Gene transcripts were identified based on their species name followed by count number. Sequences denoted with a six-digit alphanumeric code are from the Swiss-Prot/UniProt databases.

#### 3.3.3 Inhibitor Cystine Knot

Two structurally similar ICK (Kasheverov et al., 2022; Pallaghy et al., 1994) toxins were identified in the transcriptomes, a 6-cysteine (Figure 3) and 8-cysteine (Figure 4) framework. Both types have similarity to ICK toxins in spider species (such as, Uniprot accessions P61095, P60514 and D2Y1X8), but the 6-cysteine framework has the closest similarity (Norton & Pallaghy, 1998). Of note, sequences with six cysteines have the KR cleavage site preceding the first cysteine by 0-1 residue (Figure 3a), whereas sequences with eight cysteines have no cleavage site (Figure 4a), suggesting a mature peptide directly after the signal peptide. There is a potential cleavage site between the second and third cysteine (position 57-58), which requires further investigation (Figure 4a). The 8-cysteine framework has very strong conservation of residues for the mature peptide, and there is no variation in the number of residues between the first and last cysteines (X_n_-C-X_6_-C-X_4_-CC-X_1_-CC-X_4_-C- X_12_-C-X_n_) (Figure 4a). This framework has similarity to the spider toxin, Lambda-hexatoxin- Hv1c, found in *Hadronyche versuta* (UniProt Accession: P82228; Wang et al., 2000). Interestingly, six of the seven sequences in the 6-cysteine group were found in superfamily Metridioidea, whereas eight of the nine sequences in the 8-cysteine group were found in superfamily Actinioidea, suggesting independent expansions of these peptides given that the only overlap between these groups is *M. senile*. Phylogenetic analysis for the 6-cysteine group had a distinct clustering by lineage (Figure 3b), whereas the 8-cysteine group had very strong clustering except for the Ioli_1 and Avt_1 group (Figure 4b), although these two sequences only vary by one amino acid (Figure 4a).

**Figure 3:**
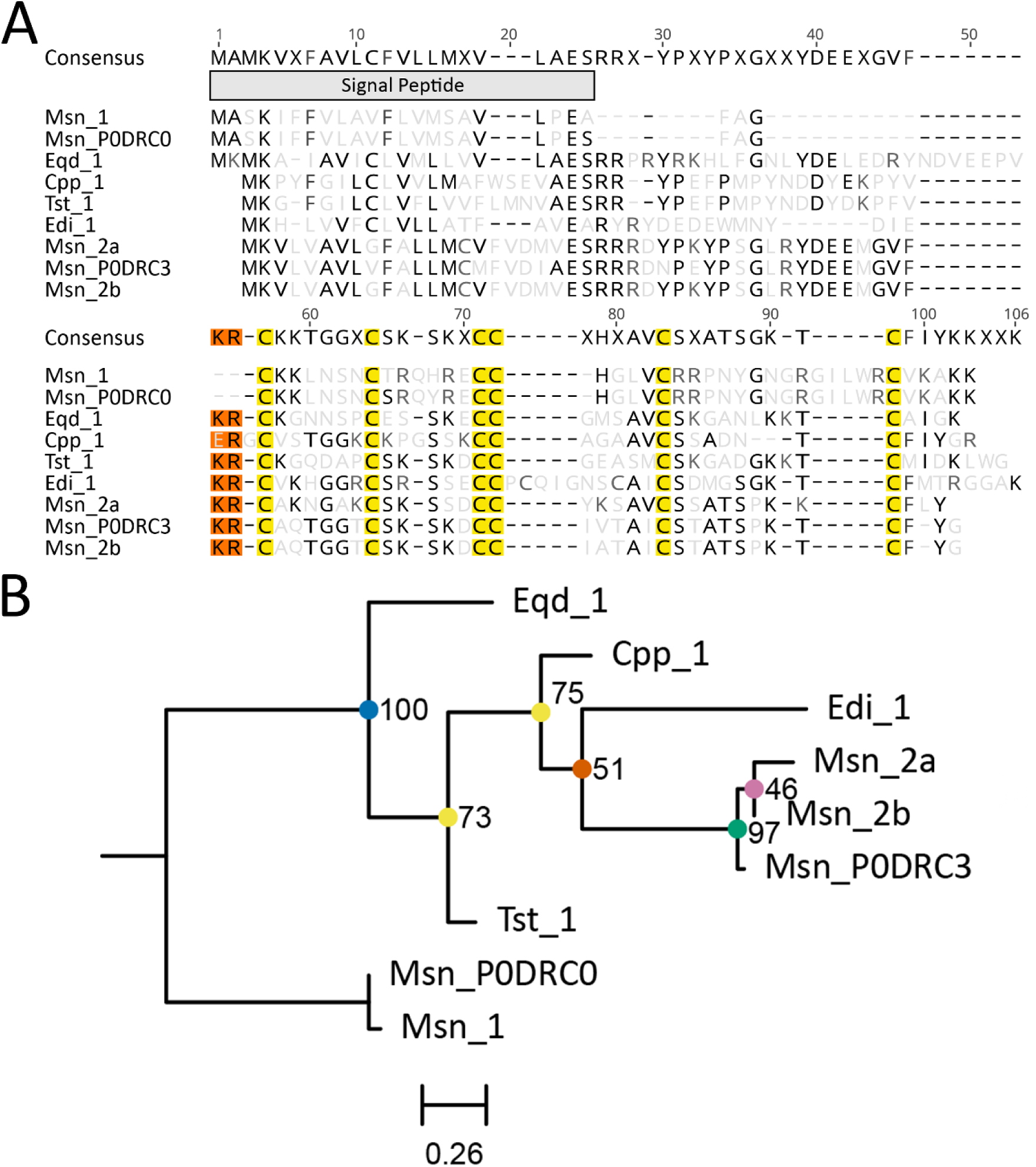
Amino acid sequence alignment and phylogenetic tree for the 6-cysteine ICK toxin family sequences. (A) Alignment of candidate sequences with highlighted sections for predicted signal peptide, cysteine residues and potential cleavage site motifs. Light grey letters represent non-consensus amino acids. (B) Maximum-likelihood phylogenetic tree of candidate toxins. Support values shown as Maximum Likelihood bootstrap (0-100). Gene transcripts were identified based on their species name followed by count number or isoform designation denoted with alphanumeric values. Sequences denoted with a six-digit alphanumeric code are from the Swiss-Prot/UniProt databases.

**Figure 4:**
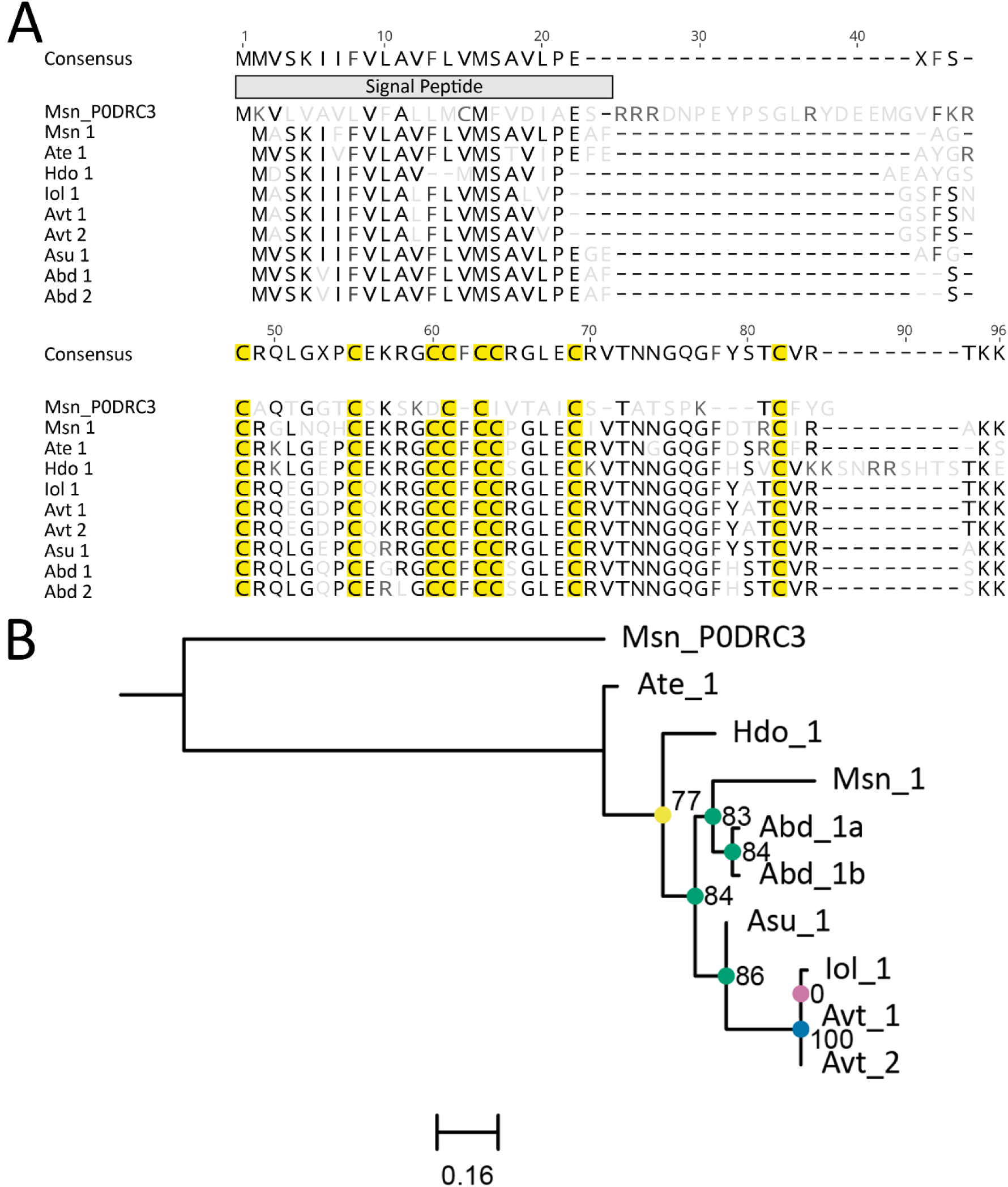
Amino acid sequence alignment and phylogenetic tree for the 8-cysteine ICK toxin family sequences. (A) Alignment of candidate sequences with highlighted sections for predicted signal peptide and cysteine residues. Light grey letters represent non-consensus amino acids. (B) Maximum-likelihood phylogenetic tree of candidate toxins. Support values shown as Maximum Likelihood bootstrap (0-100). Gene transcripts were identified based on their species name followed by count number or isoform designation denoted with alphanumeric values. Sequences denoted with a six-digit alphanumeric code are from the Swiss-Prot/UniProt databases.

#### 3.3.4 Phospholipase A2

A total of 157 PLA2 (Talvinen & Nevalainen, 2002) toxin candidates was found across 26 species of sea anemones, with three cysteine frameworks identified: 10-, 12- and 14-cysteine (Figure 5a, Figure 5b and Figure 5c, respectively) (Supplementary Figure 1, Clade 1, Clade 2 and Clade 3, respectively). The 10- and 12-cysteine frameworks have been identified previously in sea anemones (Martins et al., 2009; Razpotnik et al., 2010; Romero et al., 2010; Smith et al., 2023; Smith et al., 2024; Talvinen & Nevalainen, 2002), while the 14-cysteine framework has only been reported in other venomous animals. Expected cleavage sites for mature peptide sequences varied among the three cysteine frameworks (Table 3), with only 103 having the canonical KR motif, whereas all sequences for the 10-cysteine framework had no known cleavage motifs. Phylogenetic analyses for the entire repertoire of PLA2 toxins in sea anemones revealed one cluster for each of the three structural types, and strong support was seen for the three clades. Interestingly, there are two different consensus sequences for the 12-cysteine framework with sequence alignments having either a X_10_-C-X_15_-C-X_14_-CC-X_5_-C-X_6_-C-X_17_-C-X_3_-C-X_6_-C-X_4_-C-X_1_-C-X_6_-C-X_n_ or X_22_-C-X_1_-C-X_14_-CC-X_5_-C-X_11_-C-X_16_-C-X_8_-C-X_4_-C-X_1_-C-X_6_-C-X_17_-C-X_n_ motifs (Supplementary Figure 2).

**Figure 5:**
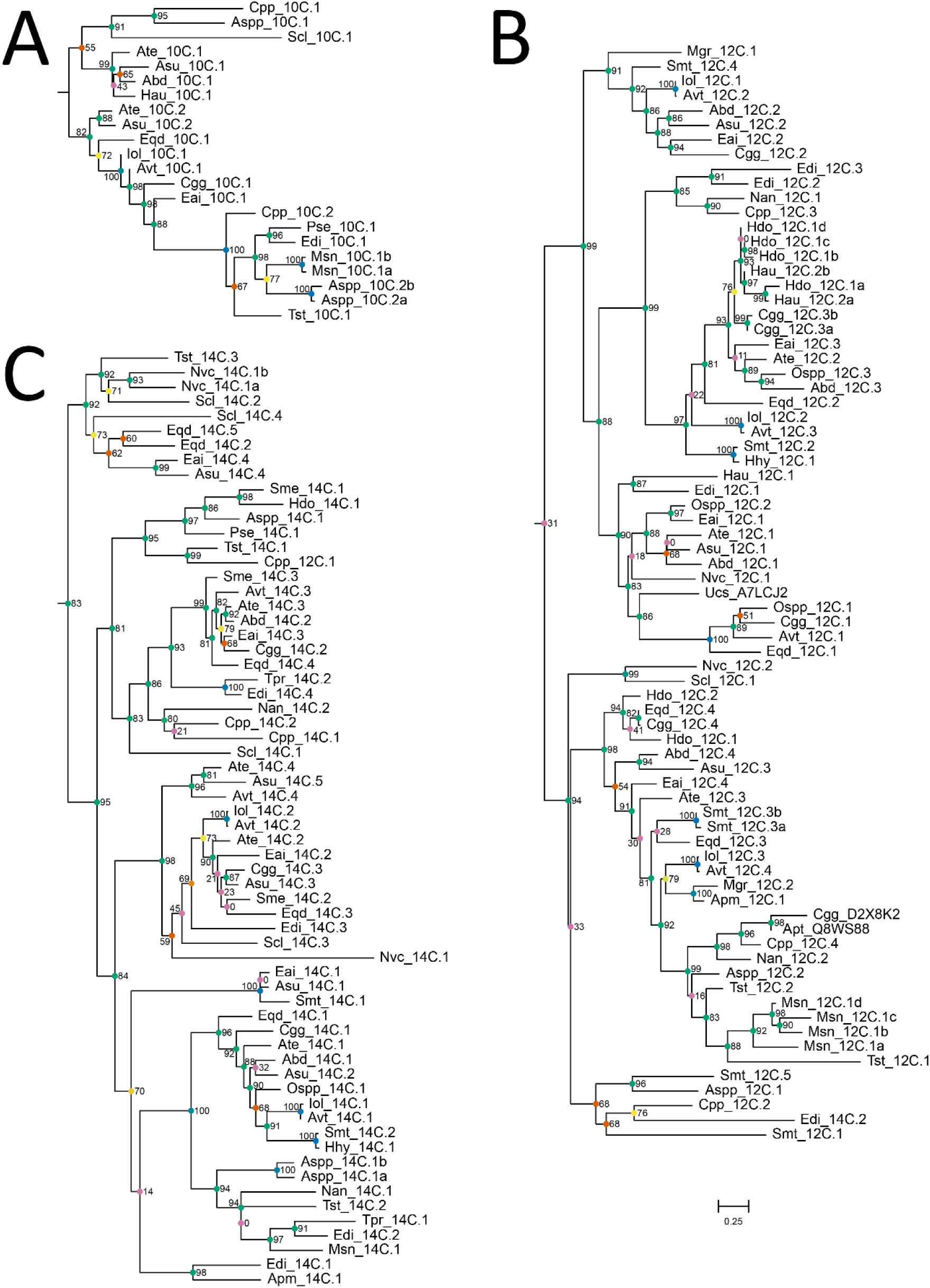
Maximum-likelihood phylogenetic tree of candidate sequences for the phospholipase A2 toxin family. A. Sequences forming the 10-cysteine framework (Clade 1 of Supplementary Figure 1). B. Sequences forming the 12-cysteine framework (Clade 2 of Supplementary Figure 1). C. Sequences forming the 14-cysteine framework (Clade 3 of Supplementary Figure 1). Support values shown as Maximum Likelihood bootstrap (0-100). Gene transcripts were identified based on their species name followed by count number or isoform designation denoted with alphanumeric values. Sequences denoted with a six-digit alphanumeric code are from the Swiss-Prot/UniProt databases. Scale bar shown in C is also applicable for A and B.

**Table 3:** Summary of potential cleavage site motifs found in phospholipase A2 sequences (n denotes an E, G, S or V residue, and N/A denotes no cleavage site)

| Cystine Framework | Cleavage Motif |  |  |  |  |
| --- | --- | --- | --- | --- | --- |
|  | KR | RR | KK | nR | N/A |
| 10C | 0 | 0 | 0 | 0 | 22 |
| 12C | 69 | 0 | 2 | 0 | 0 |
| 14C | 34 | 14 | 0 | 9 | 7 |

#### 3.3.5 Potassium channel toxin type V

Potassium channel toxin type V (Orts et al., 2013) sequences were only found in the superfamily Actinioidea, although sequences have been reported in Edwardsioidea and Metridioidea species (UniProt accessions: A7RMN1 (*N. vectensis*) and P0DMD7 (*M. senile*); Figure 6a). All candidates had potential cleavage sites KR, RR (Arg-Arg) or TR (Thr-Arg), with the exception of Smt_1, Abd 1 and Asu 1, which have mature peptides directly after the signal peptide (Figure 6a). Phylogenetic analyses had varied support values throughout the tree, although Clade 1 (Ate_1, Cgg_1 and Hau_1) and Clade 2 (Smt_1, Abd_1 and Asu_1) were strongly supported (Figure 6b). Sequences forming Clade 3 had greater similarity to known KTx type V, Msn_P0DM07, Bcs_C0HJC4 and Nvc_A7RMN1, but several internal nodes have low support values. Reanalysis of Clade 1 and Clade 2 with UniProt accessions reaffirms the separate clustering of these clades (Supplementary Figure 3) and suggests that functional characterisation of toxin candidates in species from Clade 1 and 2 is needed.

**Figure 6:**
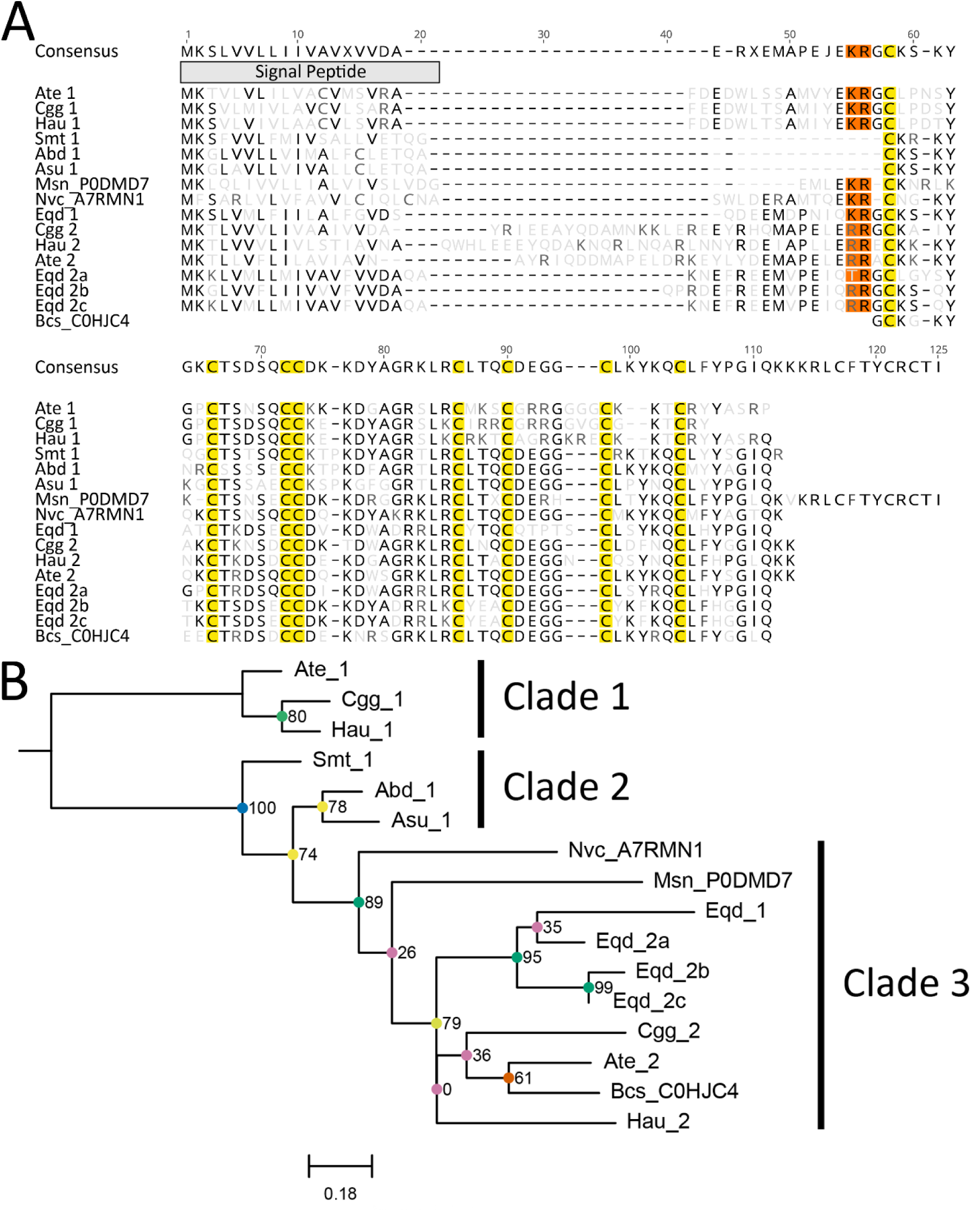
Amino acid sequence alignment and phylogenetic tree for the potassium channel toxin type V family sequences. (A) Alignment of candidate sequences with highlighted sections for predicted signal peptide, cysteine residues and potential cleavage site motifs. Light grey letters represent non-consensus amino acids. (B) Maximum-likelihood phylogenetic tree of candidate toxins. Support values shown as Maximum Likelihood bootstrap (0-100). Gene transcripts were identified based on their species name followed by count number or isoform designation denoted with alphanumeric values. Sequences denoted with a six-digit alphanumeric code are from the Swiss-Prot/UniProt databases.

#### 3.3.6 Small cysteine-rich protein

Small cysteine-rich proteins (Jouiaei et al., 2015) were found in several species across superfamilies Actinioidea and Metridioidea but were absent in superfamily Edwardsioidea. All but three sequences (Apm_1, Mgr_1b and Iol_1) and UniProt accession C1KIZ4 (*Orbicella* sp.), had a KR or RR cleavage motif (Figure 7a). In superfamily Actinioidea, three candidates, Eai_1, Mgr_1 and Eqd_1, had multiple gene isoforms: two, two and three, respectively. The Eqd_1 gene isoforms all clustered closely together with strong support for an independent clade, while all other genes, except Ospp_1, grouped with the three stony coral sequences (Figure 7b).

**Figure 7:**
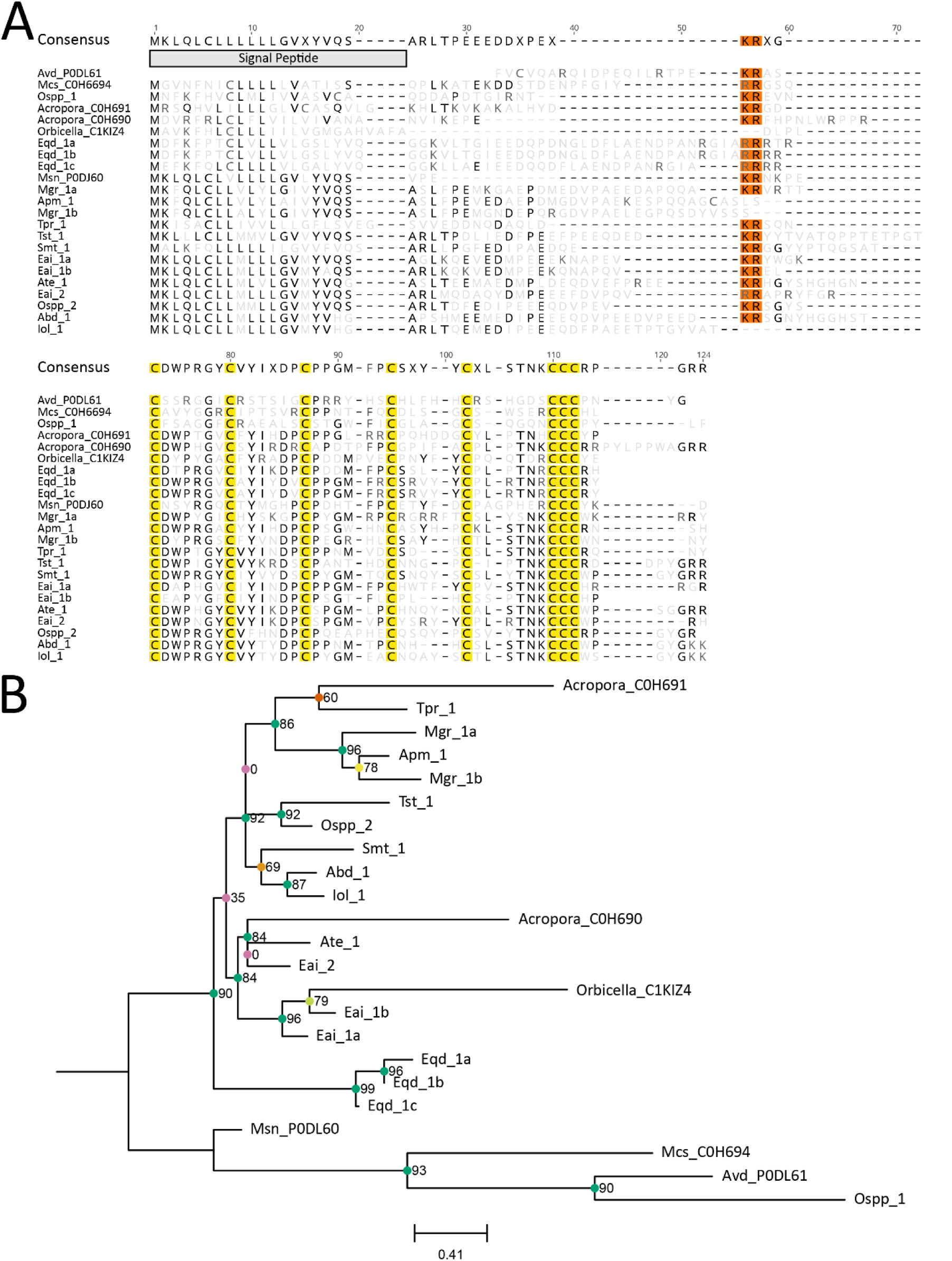
Amino acid sequence alignment and phylogenetic tree for the small cysteine-rich protein toxin family sequences. (A) Alignment of candidate sequences with highlighted sections for predicted signal peptide, cysteine residues and potential cleavage site motifs. Light grey letters represent non-consensus amino acids. (B) Maximum-likelihood phylogenetic tree of candidate toxins. Support values shown as Maximum Likelihood bootstrap (0-100). Gene transcripts were identified based on their species name followed by count number or isoform designation denoted with alphanumeric values. Sequences denoted with a six-digit alphanumeric code are from the Swiss-Prot/UniProt databases.

#### 3.3.7 Sodium channel toxin type III (Short III)

Only five NaTx type III (Manoleras & Norton, 1994; Martinez et al., 1977) candidates were identified in the 26 transcriptomes analysed, with Eqd_1 matching P09949 and Asu_1 closely matching C3TS04 (Figure 8). The three newly identified sequences, Cad_1, Hhy_1 and Ate_1, have different cysteine frameworks from these two toxins and were more similar to Asu_1 than Eqd_1, except for the number of cysteine residues. Strong support was observed for two phylogenetic clades and low support was observed for Hhy_1 and Cad_1 (Figure 8b). This coincides with the known structural variants, P09949 (*E. quadricolor*) and C3TS04 (*A. viridis*), compared to the alternative 8-cysteine framework (KR-X_2_-CC-X_1_-C-X_2_-C-X_1_-C-X_4_- CC-X_4_-C) found in Cad_1, Hhy_1 and Ate_1 (Figure 8a).

**Figure 8:**
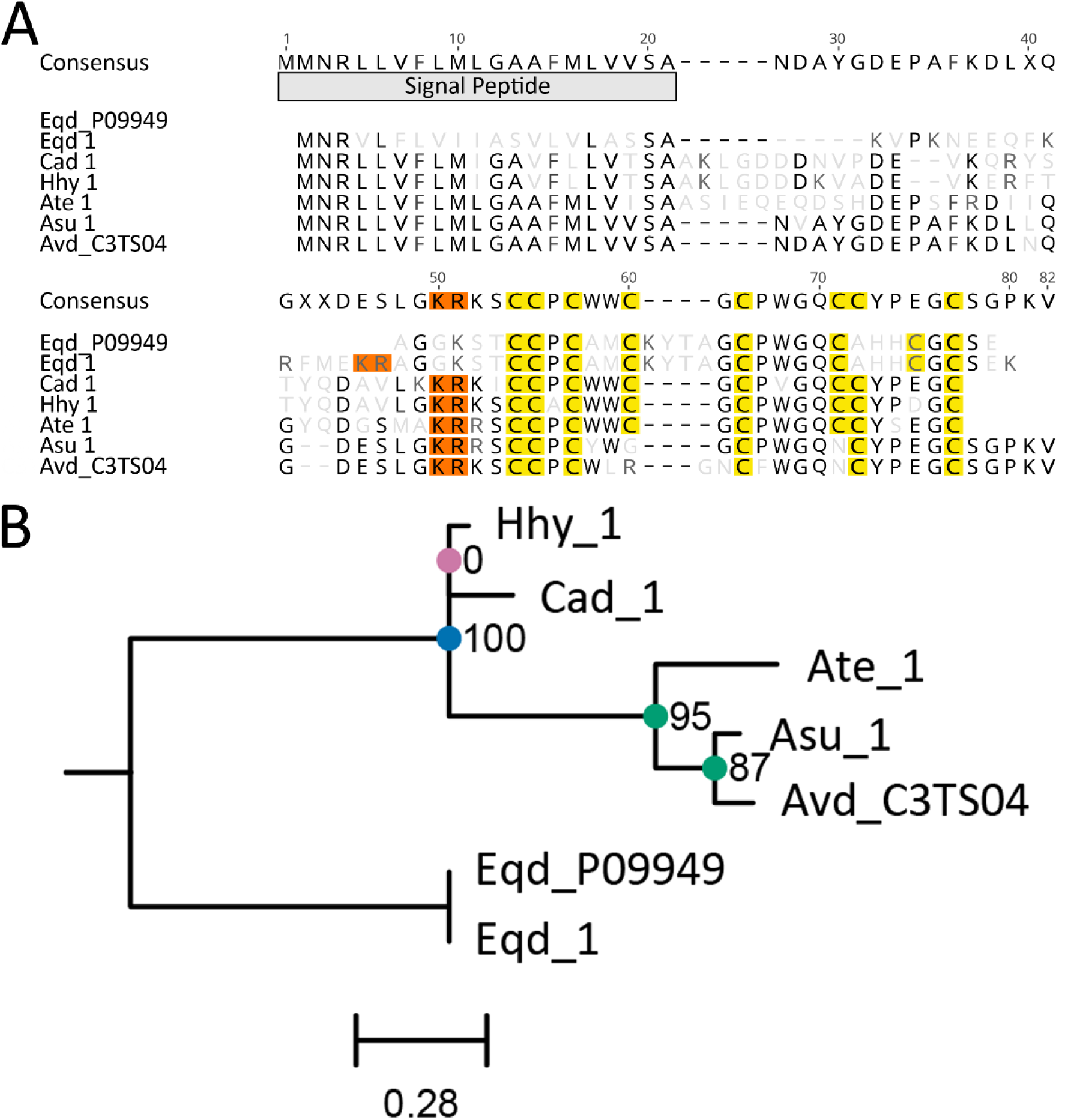
Amino acid sequence alignment and phylogenetic tree for the sodium channel toxin type III family sequences. (A) Alignment of candidate sequences with highlighted sections for predicted signal peptide and cysteine residues. Light grey letters represent non-consensus amino acids. (B) Maximum-likelihood phylogenetic tree of candidate toxins. Support values shown as Maximum Likelihood bootstrap (0-100). Gene transcripts were identified based on their species name followed by count number or isoform designation denoted with alphanumeric values. Sequences denoted with a six-digit alphanumeric code are from the Swiss-Prot/UniProt databases.

## 4. Discussion

The distribution of toxin gene families in the six new species is consistent with other species from superfamily Actinioidea, although toxin gene copy numbers varied greatly among the new species, as well as with other species from their respective superfamilies. For example, there were no copies of KTx type III identified in *A. plumosum*, even though this gene family is common in other species from superfamily Actinioidea (Macrander et al., 2016; Ramirez- Carreto et al., 2019; Surm et al., 2019b); in fact, there are 5-10 copies found in the other five species reported in this study. Similarly, copy numbers of the PLA2 family were lower in *A. plumosum*, *C. adhaesivum* and *H. aurora* compared to most species, whereas the KTx type II family had the most abundant copy numbers in *C. adhaesivum* and *S. mertensii*. This pattern contrasts with the proposed ubiquitous distribution and high copy number of these toxin families across sea anemone species that has been proposed previously (Chen et al., 2019; Macrander et al., 2016; Madio et al., 2017; Prentis et al., 2018; Surm et al., 2019b). Overall, our data suggest that toxin gene copy numbers derived from transcriptomic data are highly dynamic across most toxin families, and toxin copy numbers were surprisingly lower in the medically significant species, *A. plumosum*, when compared to other species in the same superfamily.

### 4.1 Evolution and diversity of toxin gene families in sea anemones

Among sea anemone taxa there is an irregular distribution of genes encoding toxin families. Based on the seven toxin families we analysed, genes encoding AcrI, CjTL, 8-cysteine ICK, KTx type V, NaTx type III and SCRiP toxins were more frequently identified in superfamily Actinioidea and 6-cysteine ICK toxins were predominantly found in Metridioidea, while only PLA2 was commonly found in all superfamilies. Similar patterns among sea anemone superfamilies have been observed previously for toxin gene families with restricted (Barroso et al., 2024; Surm et al., 2024b) or widespread distributions (Macrander & Daly, 2016; Surm et al., 2019a;). Many toxin-encoding genes are patchily distributed across sea anemone species, with high levels of extensive toxin gene loss and gain reported in different lineages (Jouiaei et al., 2015; Macrander & Daly, 2016; Macrander et al., 2016; Madio et al., 2018; Mitchell et al., 2019; Surm et al., 2019a; Surm et al., 2019b). Our data for these toxin families also support this pattern of gene loss and gain in the evolution of toxin gene families in sea anemones.

Further investigation of the neurotoxins AcrI, CjTL, ICK, KTx type V, NaTx type III and SCRiP, revealed a pattern of lineage-specific gene gain. A limited distribution of genes encoding these toxins was observed, with the highest number of gene copies often found in superfamily Actinioidea. The overall copy numbers for these genes were typically less than other neurotoxins that have been studied previously (Ashwood et al., 2023; Jouiaei et al., 2015; Surm et al., 2019a). For example, in our data, the sea anemone 8 toxin (Ashwood et al., 2023) had over four times the total number of genes compared with SCRiP, which had the highest gene count of the neurotoxins we analysed. The restricted occurrence of specific toxins in our data could be the result of tissue specificity and/or lineage-specific evolution. For example, AcrI was originally identified in the acrorhagi, a specialised cellular structure, of *A. equina* (Honma et al., 2005) and this structure is found in very few species of family Actiniidae. Despite this, our data suggest that an AcrI-like peptide is found in a more diverse range of species without acrorhagi, giving rise to the possibility of a third structural form (alternative to acrorhagin type I and II) that is not associated with acrorhagi. Alternatively, low copy numbers and a limited phylogenetic distribution observed in the six neurotoxins we analysed is a pattern indicative of prey/predator selectivity, a process previously reported for neurotoxic proteins found in the venom of some snake lineages (Harris et al., 2020; Michálek et al., 2024), or another unknown selective pressure is maintaining these genes.

Compared to the other toxins analysed, the PLA2 family was ubiquitously distributed across all superfamilies examined. Multiple copies of PLA2 were observed in all but one of the sea anemone species examined, with high copy number variability identified across the sea anemone families. For example, within superfamily Actinioidea, the Actinodendridae and Thalassianthidae families had 1-2 copies, whereas the Actiniidae family had 4-10 copies. The possibility of taxonomically restricted evolution in combination with gene duplication and loss likely influenced this variability, both processes having been reported for PLA2 toxins in other venomous animals (Casewell, 2016; Dowell et al., 2016). Limited analyses have been performed on PLA2 toxins in sea anemones, with only the function of the 12-cysteine protein characterised for the hydrolysis of phospholipids (Martins et al., 2009; Romero et al., 2010; Talvinen & Nevalainen, 2002) and an effect on renal function in rats (Martins et al., 2009). In other venomous animals, 10-, 12- and 14-cysteine PLA2 toxins typically have a role in prey capture/digestion and defence (Krayem & Gargouri, 2020; Sampat et al., 2023) and would likely have a similar role in sea anemones. Specifically, a 12-cysteine PLA2 protein was identified recently in the venom of *C. polypus* suggesting it may have a role in prey capture/digestion (Smith et al., 2024) and defence (Smith et al., 2023). Consequently, PLA2 toxins are an ideal candidate to study how structural variation influences toxicity and function in sea anemones as PLA2 is universally distributed and has high sequence variation.

### 4.2 The limitations of transcriptome sequences for evaluating toxin gene evolution

We observed a patchy distribution for many toxin encoding genes, a pattern that has been observed in transcriptomic studies in other venomous animals (Aird et al., 2013; Casewell et al., 2009; Jiang et al., 2011; Ma et al., 2012). In transcriptomic studies it can be difficult to determine whether genes are truly absent or are missing as a result of dynamic expression patterns or low levels of gene expression. For example, NvePTx1, a KTx type V gene found in *N. vectensis*, has restricted expression to the early life stages and adult females during egg formation (Columbus-Shenkar et al., 2018). Similarly, transcriptomic data for SCRiP toxins in stony corals suggests that some genes are differentially expressed across life stages (Barroso et al., 2024). Given that our current study relies on transcriptomic data generated from adult organisms, those genes expressed in distinct life stages may not have been captured in our analysis. One example of this disparity is the lack of any MACPF transcripts in the *A. tenebrosa* transcriptome, although two MACPF genes are present in the genome sequence of *A. tenebrosa* (Surm et al., 2019a; Surm et al., 2024b). This indicates that we have not captured the full complexity of the MACPF gene family in our transcriptomic data and subsequently the whole toxin complement. Future studies examining toxin gene complements in non-model sea anemones would greatly benefit from the use of a combination of venom proteome, transcriptome (including differential life stages), and genome assemblies to accurately identify all gene copies in toxin families. Such an approach will allow a more precise understanding of the molecular evolution of toxins and may also overcome issues with ascertainment bias when using transcriptomic data in isolation.

The low toxin copy number reported in the *A. plumosum* transcriptome assembly may result from ascertainment bias and limitations of current toxin characterisations, particularly as it had the largest number of transcripts in this study. The majority of previously published work has characterised toxins from a limited diversity of sea anemone families, such as Actiniidae, Edwardsiidae, Heteractidae, and Stichodactylidae (Columbus-Shenkar et al., 2018; Honma et al., 2005; Honma et al., 2008; Macrander et al., 2016; Madio et al., 2017; Mitchell et al., 2020; Moran et al., 2008; Moran et al., 2013; Rivera-de-Torre et al., 2018; Sachkova et al., 2019; Surm et al., 2019b), which are distantly related to two of the six species that we analysed, *A. plumosum* and *C. adhaesivum* (Rodríguez et al., 2014). Thus, it is not surprising we found low copy numbers in these two species – as the venom components of species from Actinodendridae and Thalassianthidae have not been examined in detail, and many undiscovered toxin peptides and proteins may exist in these families. A combined proteo- genomic approach, such as that employed by others (Ashwood et al., 2022; Madio et al., 2017; Ramirez-Carreto et al., 2019), may have yielded a greater diversity of toxins as this approach has been shown to identify new toxins and toxin candidates. Alternatively, it is possible that the modulation of venom in sea anemones is highly complex (Schendel et al., 2019). For example, the toxin repertoire of acontia venom has low complexity but a single neurotoxin was more highly abundant in the venom than all other toxins present (Smith et al., 2023). Therefore, determining the toxin peptide and protein abundances in sea anemone venom may elucidate possible roles of venom modulation and would likely mitigate ascertainment bias by identifying new toxins and toxin families in more diverse species, as well as for medically significant species, such as *A. plumosum*.

## 5. Conclusion

Genomic resources are helping to generate new insights in our understanding of the diversity and complexity of sea anemone venom and toxin evolution. Additional transcriptome sequencing (presented herein) allowed us to expand and investigate the evolution of previously understudied toxin gene families. Here we report that six neurotoxin families in sea anemones have undergone a process of taxonomically restricted evolution, whereas the PLA2 family is a more widely distributed toxin with high variability among taxonomic families. Our analyses encompass a taxonomically diverse group of sea anemones which provide a comprehensive list of novel toxins to establish the foundational experiments for identifying candidates for future toxinological analyses. Elucidating the structure and function of these novel toxins would further our understanding of their role in sea anemones and would greatly benefit both the agricultural and therapeutic applications of these toxins.

## Data Access

All raw reads were submitted to the NCBI Sequence Read Archive (SRA) under BioProject PRJNA507679.

## CRediT authorship contribution statement

**HLS**: Conceptualization, Data Curation, Formal Analysis, Investigation, Methodology, Software, Validation, Visualization, Writing – original draft, review and editing; **DAB**: Conceptualization, Supervision, Writing – review and editing; **CAV**: Investigation, Writing – review and editing; **JMS**: Investigation, Writing – review and editing; **LL**: Investigation, Writing – review and editing; **RSN**: Writing – review and editing; **PJP**: Conceptualization, Methodology, Resources, Supervision, Writing – review and editing.

## Acknowledgements

This work was supported by the Central Analytical Research Facility, Queensland University of Technology. Computational and data visualisation resources and services used in this work were provided by the eResearch Office, Queensland University of Technology. Some of these samples and derived data have Disclosure Notices (Attribution Incomplete and Biocultural Notice) attached that are visible in a Local Contexts Hub project (9b8da5f7-8290-4547-9a4b- 36dc8ccc6d57). These Notices acknowledge that there is incomplete, inaccurate, or missing attribution, and in recognition of the rights of Indigenous peoples to define the use of these data generated from the specimens associated with their lands and waters. The Biocultural Notice indicates that Biocultural Labels may be in development. For more information, visit https://localcontextshub.org/.

