## Supplementary Figure 1-3 for "A comparative analysis of toxin gene families across diverse sea anemone species"

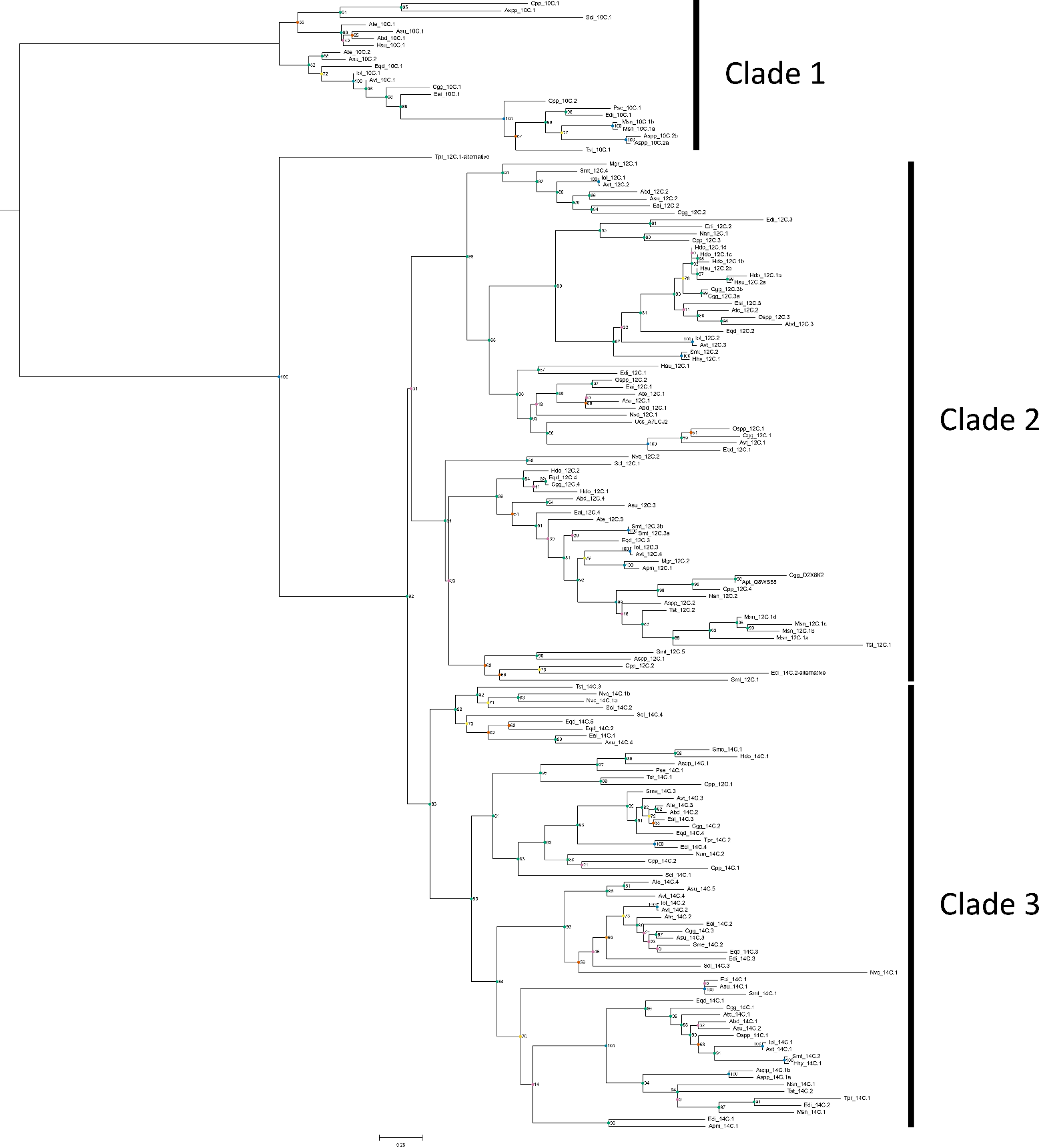

Supplementary Figure 1: Maximum-likelihood phylogenetic tree of candidate sequences for the phospholipase A2 toxin family. Support values shown as Maximum Likelihood bootstrap (0-100). Gene transcripts were identified based on their species name followed by count number or isoform designation denoted with alphanumeric values. Sequences denoted with a six-digit alphanumeric code are from the Swiss-Prot/UniProt databases.

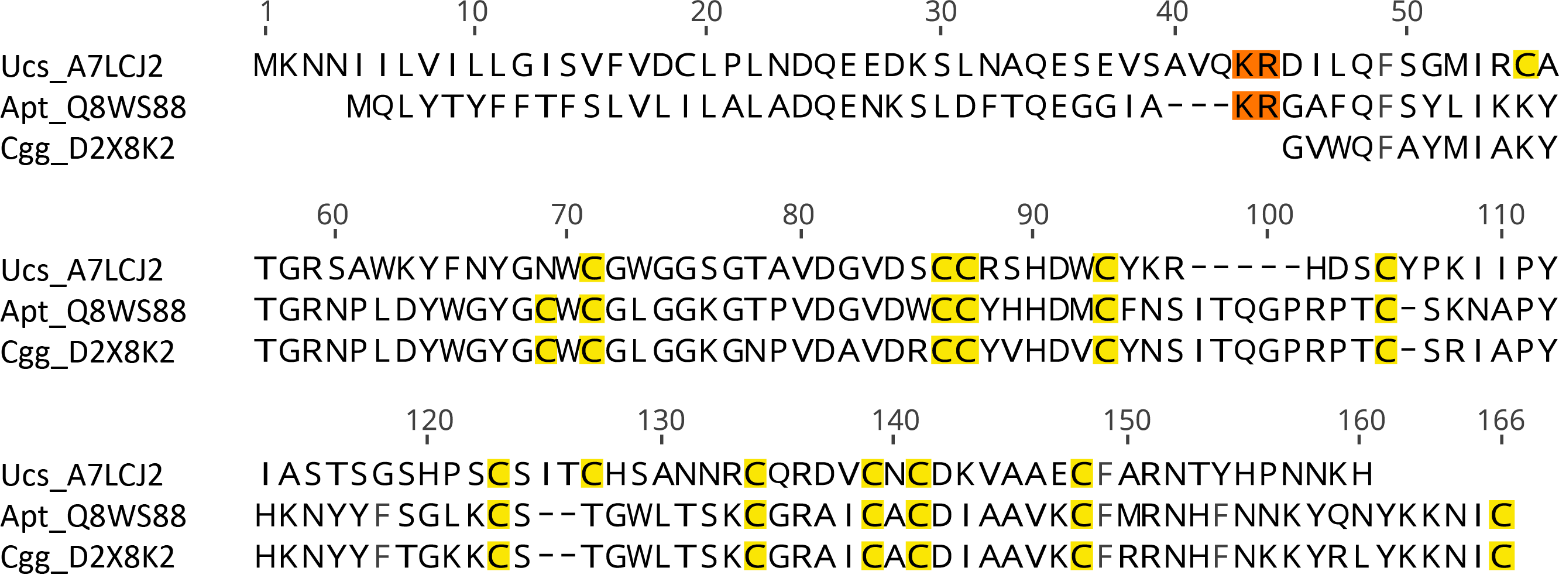

Supplementary Figure 2: Alignment of characterised 12-cysteine phospholipase A2 sequences from UniProt showing two cysteine framework structures, with highlighted sections for cysteine (C) residues and known cleavage site motifs (KR). Abbreviations: *Urticina crassicornis* (Ucs), *Calliactis palliata* (Apt; formerly *Adamsia palliata*), *Condylactis gigantea* (Cgg). Sequences denoted with a six-digit alphanumeric code from the Swiss-Prot/UniProt databases.

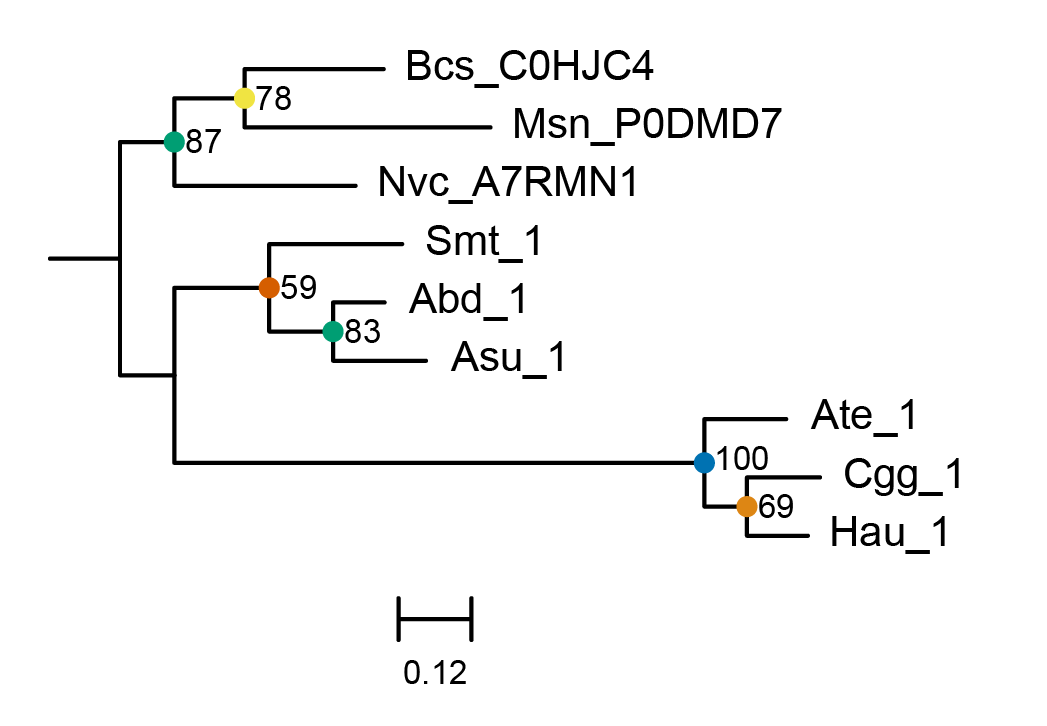

Supplementary Figure 3: Maximum-likelihood phylogenetic tree for the potassium channel toxin type V family sequences derived from Clade 1 and Clade 2 of Figure 6 in Manuscript. Support values shown as Maximum Likelihood bootstrap (0-100). Gene transcripts were identified based on their species name followed by count number or isoform designation denoted with alphanumeric values. Sequences denoted with a six-digit alphanumeric code are from the Swiss-Prot/UniProt databases.
